# Recombination-driven genome evolution and stability of bacterial species

**DOI:** 10.1101/067942

**Authors:** Purushottam D. Dixit, Tin Yau Pang, Sergei Maslov

## Abstract

While bacteria divide clonally, horizontal gene transfer followed by homologous recombination is now recognized as an important and sometimes even dominant contributor to their evolution. However, the details of how the competition between clonal inheritance and recombination shapes genome diversity, population structure, and species stability remains poorly understood. Using a computational model, we find two principal regimes in bacterial evolution and identify two composite parameters that dictate the evolutionary fate of bacterial species. In the divergent regime, characterized by either a low recombination frequency or strict barriers to recombination, cohesion due to recombination is not sufficient to overcome the mutational drift. As a consequence, the divergence between any pair of genomes in the population steadily increases in the course of their evolution. The species as a whole lacks genetic coherence with sexually isolated clonal sub-populations continuously formed and dissolved. In contrast, in the metastable regime, characterized by a high recombination frequency combined with low barriers to recombination, genomes continuously recombine with the rest of the population. The population remains genetically cohesive and stable over time. The transition between these two regimes can be affected by relatively small changes in evolutionary parameters. Using the Multi Locus Sequence Typing (MLST) data we classify a number of well-studied bacterial species to be either the divergent or the metastable type. Generalizations of our framework to include fitness and selection, ecologically structured populations, and horizontal gene transfer of non-homologous regions are discussed.

## INTRODUCTION

Bacterial genomes are extremely variable, comprising both a consensus ‘core’ genome which is present in the majority of strains in a population, and an ‘auxiliary, genome, comprising genes that are shared by some but not all strains (1-7).

Multiple factors shape the diversification of the core genome. For example, random point mutations generate single nucleotide polymorphisms (SNPs) within the population that are passed on *vertically* from mother to daughter. At the same time, stochastic elimination of lineages leads to random fixation of polymorphisms which effectively reduces population diversity. The balance between point mutations and random fixation determines the average number of genetic differences between pairs of individuals in a population, often denoted by *θ.*

During the last two decades, exchange of genetic fragments between closely related organisms has also been recognized as a significant factor in bacterial evolution (5, 6, 8–13). Transferred genetic segments are integrated into the recipient chromosome via homologous recombination. Notably recombination between pairs of strains is limited by the divergence in transferred regions.

The probability *p*_success_ *∼*e^-*δ*/*δ*^_TE_ of successful recombination of foreign DNA into a recipient genome decays exponentially with *δ*, the local divergence between the donor DNA fragment and the corresponding DNA on the recipient chromosome (11, 14–17). In this work, we refer to *δ*_TE_ as the transfer efficiency. *δ* _TE_ is shaped at least in part by the restriction modification (RM), the mismatch repair (MMR) systems, and the biophysical mechanisms of homologous recombination (14, 15). The transfer efficiency *δ*_TE_ imposes an effective limit on the divergence among subpopulations that can successfully exchange genetic material with each other (14, 15).

On the one hand, vertical inheritance of polymorphisms leads to a clonally structured population wherein genomes of mothers and daughters are very similar to each other. On the other hand, recombinations of genetic fragments within the population can exchange polymorphisms horizontally, potentially destroying the genetic signatures of clonal relationships (6, 16–18). As a result of the competition between these two factors, bacterial genomes can have varying degree of clonality.

Computational studies have explored some aspects of this competition. For example, Falush et al. (19) suggested that a low transfer efficiency *δ*_TE_ leads to sexual isolation in *Salmonella enterica;* strains within individual: subclades exchange genes among themselves but rarely: between clades. In contrast, Fraser et al. (16), working: with *θ* = 0.4% (lower than typical *θ*s in real bacterial: populations) and the transfer efficiency *δ*_TE_ ≈ 2.4% concluded that realistic recombination rates are insufficient to cause sexual isolation. In a more general study, Doroghazi and Buckley (20), working with a fixed transfer efficiency and a very small population size (limit of *θ* → 0: of our study), studied the parameter ranges where sexual isolation between subclades can be overcome through a combination of high recombination rates and/or high transfer efficiency. However, a clear understanding of the competition between recombinations and mutations remains over a broad range of evolutionary parameters remains elusive.

In this work, we develop an evolutionary theoretical framework that allows us to study in broad detail the nature of competition between recombinations and point mutations. We identify two composite parameters that: govern how genes and genomes diverge from each other over time. Each of the two parameters corresponds to: a competition between vertical inheritance of polymorphisms and their horizontal exchange via homologous recombination.

First is the competition between the recombination rate *ρ* and the mutation rate *μ.* The recombination rate: *ρ* depends on the mechanisms (21) available for genetic exchange including transformation, conjugation, transduction, etc. Within a co-evolving population, consider: a pair of strains diverging from each other. The average time between consecutive recombination events affecting: any given small genomic region in these two strains is 1/(2 ρ*l*_*tr*_) where *l*_*tr*_ is the average length of transferred: regions. At the same time, the total divergence accumulated in this region due to mutations in either of the two genomes is *δ*_mut_ *∼*2μ/2ρ*l*_*tr*_. If *δ*_mut_ ≫ *δ*_TE_, the pair of genomes is likely to become sexually isolated from each other in this region within the time time that separates two successive recombination events. In contrast, if *δ*_mut_ < *δ*_TE_, frequent recombination events would delay sexual isolation resulting in a more homogeneous population.

Second is the competition between the diversity within the population *θ* and the effective divergence limit *δ*_TE_ within a single sub-population uniformly capable of successful recombinations. Note that *θ* is the average pair-: wise divergence between the transferred segment and the corresponding segment on the recipient genome. If *δ*_TE_≪*θ*, one expects spontaneous fragmentation of the: entire population into several transient sexually isolated sub-populations that rarely exchange genetic material between each other. In contrast, if *δ*_TE_≫ *θ*, unhindered: exchange of genetic fragments may result in a single cohesive population.

In this work, using a computational model and mathematical calculations, we show that the two composite parameters identified above, *θ/δ*_TE_ and *δ*_mut_/*δ*_TE_, determine qualitative evolutionary dynamics of bacterial species. Furthermore, we identify two principal regimes of this dynamics. In the divergent regime, characterized by a high *δ*_mut_/*δ*_TE_, local genomic regions acquire multiple mutations between successive recombination events and rapidly isolate themselves from the rest of the population. The population remains mostly clonal where transient sexually isolated sub-populations are continuously formed and dissolved. In contrast, in the metastable regime, characterized by a low *δ*_mut_/*δ*_TE_ and a low *θ*/*δ*_TE_), local genomic regions recombine repeatedly before ultimately escaping the pull of recombination (hence the name “metastable”). At the population level, in this regime all genomes can exchange genes with each other resulting in a genetically cohesive and temporally stable population. Notably, our analysis suggests that only a small change in evolutionary parameters can have a substantial effect on evolutionary fate of bacterial genomes and populations.

We also show how to classify bacterial species using the conventional measure of the relative strength of recombination over mutations, *r*/*m* (defined as the ratio of the number of single nucleotide polymorphisms (SNPs) brought by recombinations and those generated by point mutations in a pair of closely related strains), and our second composite parameter *θ*/*δ*_TE_. Based on our analysis of the existing MLST data, we find that different real-life bacterial species belong to either divergent or metastable regimes. We discuss possible molecular mechanisms and evolutionary forces that decide the role of recombination in a species’ evolutionary fate. We also discuss possible extensions of our analysis to include adaptive evolution, effects of ecological niches, and genome modifications such as insertions, deletions, and inversions.

## RESULTS

### Computational model

We consider a population of *N*_*e*_ co-evolving bacterial strains. The population evolves with non-overlapping generations and in each new generation each of the strains randomly chooses its parent (22). As a result, the population remains constant over time. Strain genomes have *G* = 1000 indivisible and non-overlapping transferable units. For simplicity, in what follows we refer to these units as *genes* but note that while the average protein-coding gene in bacteria is about ∼1000 base pairs (bp) long, genomes in our simulations exchange segments of length *l*_tr_ = 5000 bp mimicking genetic transfers longer than individual protein-coding genes (6, 9). These genes acquire point mutations at a rate μ per gene per generation and recombinations into a recipient genome from a randomly selected donor genome in the population are attempted at a rate *ρ* per gene per generation. The mutations and recombination events are assumed to have no fitness effects (later on we discuss how this assumption can be relaxed). In the absence of recombination *(ρ* = 0), the pairwise diversity within this population is given by *θ* = *2μN*_*e*_ (22). Finally, the probability of a successful integration of a donor gene decays exponentially, *P*_*success*_ *∼e*^-*δ*/*δ*_TE_^, with the local divergence *δ* between the donor and the recipient. Table I lists all important parameters in our model and Fig. 1 shows a cartoon illustration.

**FIG. 1.**
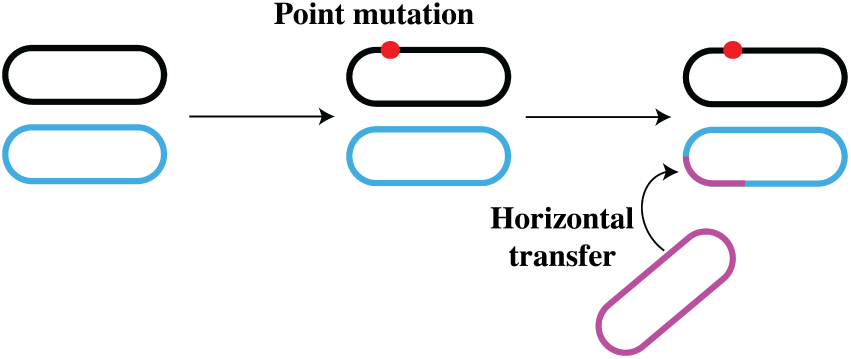
Illustration of the numerical model. *N*_*e*_ bacterial organisms evolve together, we show only one pair of strains. Point mutations (red circles) occur at a fixed rate *μ* per gene per generation and genetic fragments of length *I*_tr_ are transferred between organisms at a rate *ρ* per gene per generation.

We note that gene transfer events in bacteria may have variable end points and lengths (6). While our simplifying assumption allows us to study evolution of genome diversity extensively over a wide range of parameters, below in the discussion section, we show that our chief conclusions remain unchanged even when we relax this assumptions.

The population sizes for real bacteria are usually large (23). This prohibits simulations with realistic parameters wherein genomes of individual bacterial strains are explicitly represented. In what follows we overcome this limitation by employing an approach we had proposed earlier (6). It allows us to simulate the evolutionary dynamics of only two genomes (labeled *X* and *Y*), while representing the rest of the population using evolutionary theory (6). *X* and *Y* start diverging from each other as identical twins at time *t* = 0 (when their mother divides). We denote by *δ*_*i*_(*t*), the sequence divergences of the *i*^*th*^ transferable unit (or gene) between *X* and *Y* at time *t* and by Δ(*t*) = 1/*G* Σ_*i*_ *δ*_*i*_(*t*) the genome-wides divergence.

Based on population-genetic and biophysical conside ations, we derive the transition probability *E*(*δ*_*a*_|*δ*_*b*_) 2*μM*(*δ*_*a*_|*δ*_*b*_)+2*ρl*_*tr*_*R* (*δ*_*a*_|*δ* _*b*_) *(a* for after and *b* for before) that the divergence in any gene changes from *δ*_*b*_ to *δ*_*a*_ in one generation (6). There are two components to thes probability, *M* and *R.* Point mutations in either of twos strains, represented by *μ*(*δ*_*a*_|*δ*_*b*_), occur at a rate 2*μ* pers base pair per generation and increase the divergence in a gene by 1/*l*_*tr*_. Hence when *δ*_*a*_≠ *δ*_*b*_,

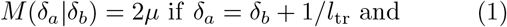

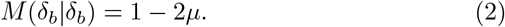

**FIG. 2.**
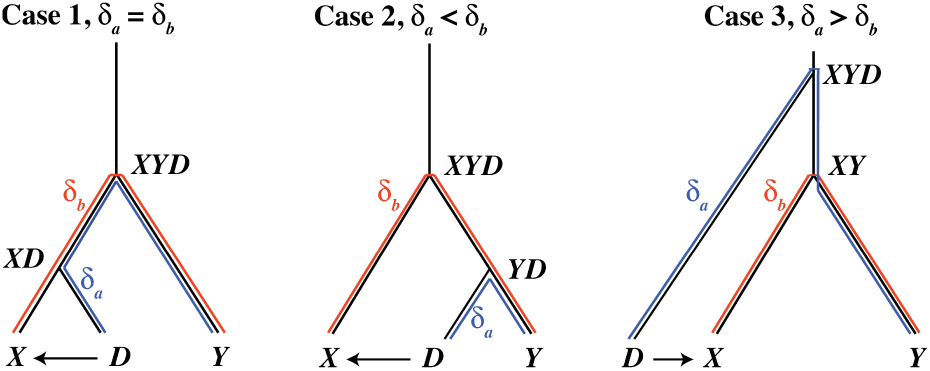
Three possible outcomes of gene transfer that change the divergence *δ. XD, YD, XY*, and *XYD* are the most recent common ancestors of the strains. The divergence *δ*_*b*_ before transfer and *δ*_*a*_ after transfer are shown in red and blue respectively.

We assume, without loss of generality, that recombination from a randomly chosen donor strain *D* within the co-evolving population replaces a gene on strain *X*. Unlike point mutations, after a recombination, local divergence between *X* and *Y* can change suddenly, taking values either larger or smaller than the current divergence (see Fig. 2 for an illustration) (6). Note that recombinations from highly diverged members in the population are suppressed exponentially and consequently not all recombination attempts are successful. We have the probabilities *R*(*δ*_*a*_|*δ*_*b*_) (6),

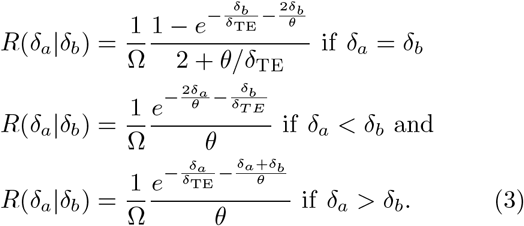

In Eqs. 3, Ω is the normalization constant.

### Evolution of local divergence has large fluctuations

Fig. 3 shows a typical stochastic evolutionary trajectory of the local divergence *δ*(*t*) of a single gene in a pair of genomes. The stochastic dynamics is simualated using the transition probability matrix *E(δ*_*a*_*|δ*_*b*_). We have used *θ* = 1.5% and *δ*_TE_ = 1%. These divergences are similar to those typically observed in bacterial species (6, 16). Mutation and recombination rates (per generation) in real bacteria are extremely small (6). In order to keep the simulation times manageable, mutation and recombination rates used in our simulations were 4-5 orders of magnitude higher compared to those observed in real bacteria (*μ* = 5 × 10^−2^ per gene per generation and *ρ* = 2.5 × 10^−2^ per gene per generation, *δ*_mut_/*δ*_TE_ = 0.04) (24, 25) while keeping the ratio of the rates realistic (5, 6, 12, 26). Alternatively, one may interpret it as one time step in our simulations being considerably longer than a single bacterial generation.

**TABLE I.** A list of parameters in the model. The range of values used in this study are indicated in the parentheses.

| parameter | symbol |
| --- | --- |
| population diversity | $\theta = 2\mu N_e$ (0.1% – 3.16%) |
| mutation rate | $\mu$ ($10^{-2}$ per gene per generation) |
| recombination rate | $\rho$ ($10^{-5} - 10^{-1}$ per gene per generation) |
| transfer efficiency | $\delta_{TE}$ (1%) |
| length of transferred regions | $l_{tr}$ (5000 base pairs) |
| number of transferable units | $G$ (1000) |

**FIG. 3.**
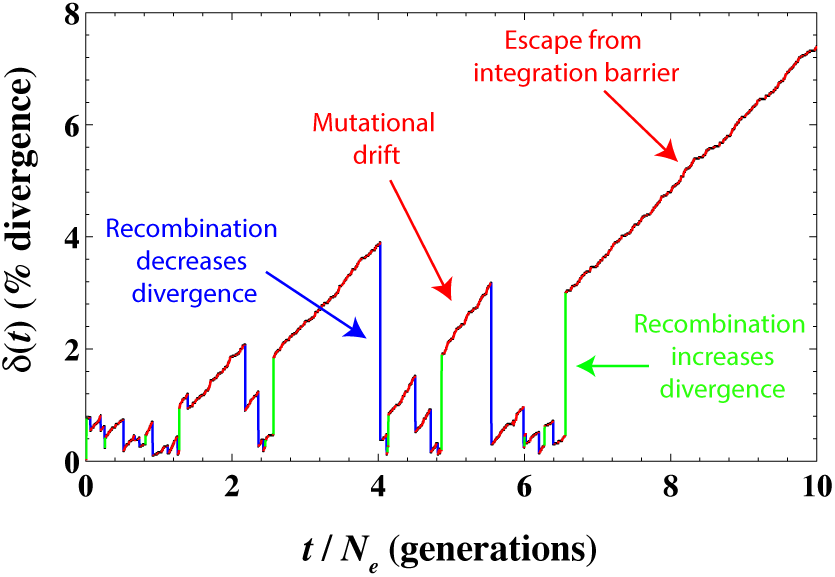
A typical evolutionary trajectory of the local divergence *δ*(*t*) within a single gene between a pair of strains. We have used *μ* = 5 × 10^−2^, *ρ* = 2.5 × 10^−2^ per gene per generation, *θ* = 1.5% and *δ*_TE_ = 1%. Red tracks indicate the divergence increasing linearly, at a rate 2*μ* per gene per generation, with time due to mutational drift. Green tracks_2_ indicate recombination events that suddenly increase the divergence and blue tracks indicate recombination events that suddenly decrease the divergence. Eventually, the divergence increases sufficiently and the local genomic region escapes the pull of recombination (red stretch at the right).

As seen in Fig. 3, the time evolution of *δ*(*t*) is noisy; mutational drift events that gradually increase the divergence linearly with time (red) are frequently interspersed with homologous recombination events (green if they increase *δ*(*t*) and blue if they decrease it) that suddenly change the divergence to typical values seen in the population (see Eq. 3). Eventually, either through the gradual mutational drift or a sudden recombination event, *δ*(*t*) increases beyond the integration barrier set by the transfert efficiency, *δ*(*t*)≫*δ*_TE_. Beyond this point, this particular gene in our two strains splits into two different sexually isolated sub-clades. Any further recombination events in this region in each of two strains would be limited to their3 sub-clades and thus would not further change the averages divergence within this gene. Conversely, the mutational drift in this region will continue to drive the two strains further apart indefinitely.

### Genome-wide divergence

Since genes in our model evolve independently of each other, the genome-wide average divergence Δ(*t*) can be calculated as the mean of *G* independent realizations of the local divergences *δ*(*t*). Since the number *G*≫1 of genes in the genome is large, the law of large numbers implies that the fluctuations in the dynamics of Δ(*t*) are substantially suppressed compared to a more noisy time course of *δ*(*t*) seen in Fig. 3.

In Fig. 4, we plot the time evolution of Δ(*t*) and its ensemble average 〈Δ(*t*)〉 (as % difference). We have used *θ* = 0.25%, *δ*_TE_ = 1%, and *δ*_mut_/*δ*_TE_ = 2, 0.2,0.04, and 2 × 10^−3^ respectively. As seen in Fig. 4, when *δ*_mut_/*δ*_TE_ is large, in any local genomic region, multiple mutations are acquired between two successive recombination events. Consequently, individual genes escape the pull of recombination rapidly and 〈Δ(*t*)〉 increases roughly linearly with time at a rate 2*μ*. For smaller values of *δ*_mut_/*δ*_TE_, the rate of change of 〈Δ(*t*)〉 in the long term decreases as many of the individual genes repeatedly recombine with the population. However, even then the fraction of genes that have escaped the integration barrier slowly increases over time, leading to a linear increase in 〈Δ(*t*)〉 with time albeit with a slope different than 2*μ*. Thus, the repeated resetting of individual *δ* (*t*)s after homologous recombination (see Fig. 3) generally results in a 〈Δ(*t*)〉 that increases linearly (albeit extremely slowly) with time.

At the shorter time scale, the trends in genome divergence are opposite to those at the longer time scale. At a fixed *θ*, a low value of *δ*_mut_/*δ*_TE_ implies faster divergence and vice versa. When recombination rate is high, genomes of strains quickly ‘equilibrate’ with the population and the genome-wide average divergence between a pair of strains reaches the population average diversity ∼*θ* (see the red trajectory in Fig. 4). From here, any new mutations that increase the divergence are constantly wiped out through repeated recombination events with the population.

Computational algorithms that build phylogenetic trees from multiple sequence alignments often rely on the assumption that the sequence divergence, for example between a pair of strains (at the level of individual genes or at the level of genomes), faithfully represents the time. that has elapsed since their Most Recent Common Ancestor (MRCA). However, Fig. 3 and Fig. 4 serve as a. cautionary tale. Notably, after just a single recombination event the local divergence at the level of individual, genes does not at all reflect time elapsed since divergence. but rather depends on statistics of divergence within a recombining population (see (6) for more details). At the. level of genomes, when *δ*_mut_/*δ*_TE_ is large (e.g. the blue. trajectory in Fig. 4), the time since MRCA of two strains. is directly correlated with the number of mutations that separate their genomes. In contrast, when *δ*_mut_/*δ*_TE_ is small (see pink and red trajectories in Fig. 4), frequent recombination events repeatedly erase the memory of the clonal ancestry. Nonetheless, individual genomic regions slowly escape the pull of recombination at a fixed rate. Thus, the time since MRCA is reflected not in the to-, tal divergence between the two genomes but in the fraction of the length of the total genomes that has escaped the pull of recombination. One will have to use a very different rate of accumulation of divergence to estimate evolutionary time from genome-wide average divergence.

**FIG. 4.**
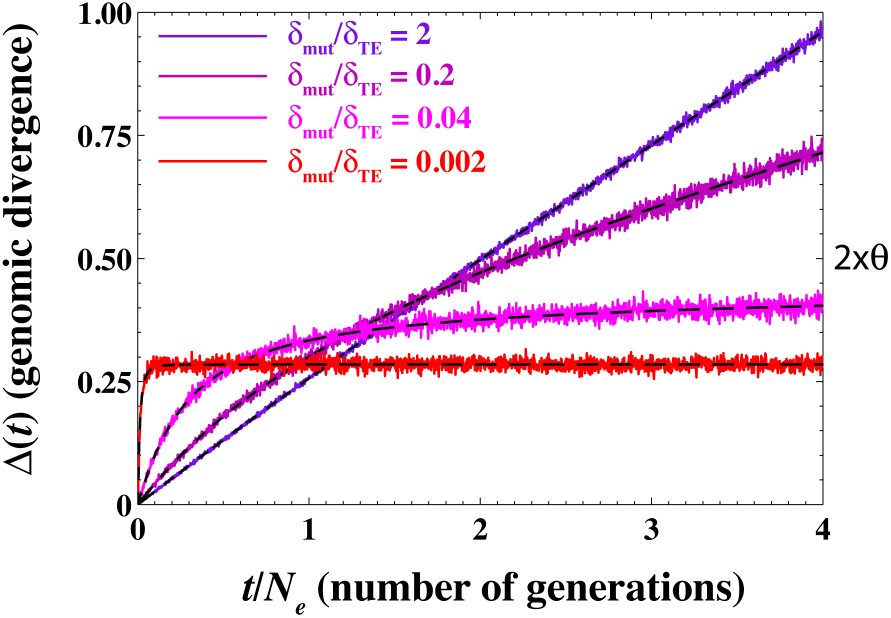
Genome-wide divergence Δ(*t*) as a function of time at *θ*/*δ*_TE_ = 0.25. We have used *δ*_TE_ = 1%, *θ* = 0.25%, μ = 10^−2^ per gene per generation and *ρ* = 10^−4^,10^−3^, 5 × 10^−2^, and 0.1 per gene per generation corresponding to *δ*_mut_/*δ*_TE_ = 2, 0.2, 0.04 and 2 × 10^−3^ respectively. The dashed black lines represent the ensemble average 〈Δ(*t*)〉. See Fig. A1 in the: appendix for the evolution of Δ(*t*) over a longer time scale.

### Quantifying metastability

How does one quantify the metastable behavior described above? At the level of individual genes it is manifested through constant resetting of *δ*(*t*) to typical population values and at the level of entire genomes through a very slow increase in Δ(*t*) when *δ*_mut_/*δ*_TE_ is small. Fig. 4 suggests that high rates of recombination prevent pairwise divergence from increasing beyond the typical population divergence ∼*θ* at the whole-genome level. Thus, for any set of evolutionary parameters, *μ*, *ρ, θ*, and *δ*_TE_, the time it takes for a pair of genomes to diverge far beyond the typical population diversity *θ* can serve as a quantifier for metastability.

In Fig. 5, we plot the number of generations *t*_div_ (in units of the effective population size *N*_*e*_) required for the ensemble average of the genome-wide average divergence 〈Δ(*t*)〉 between a pair of genomes to exceed 2 × *θ* (twice the typical intra-population diversity) as a function of *θ*/*δ*_TE_ and *δ*_mut_/*δ*_TE_. Analyzing the ensemble average 〈Δ(*t*)〉 (represented by dashed lines in Fig. 4) allows us to avoid the confounding effects of small fluctuations in the stochastic time evolution of Δ(*t*) around this average. Note that in the absence of recombination, it takes *t*_div_ = 2*N*_*e*_ generations before 〈Δ(*t*)〉 exceeds 2*θ* = 4*μN*_*e*_. The four cases explored in Fig. 4 are marked with green diamonds in Fig. 5.

We observe two distinct regimes in the behavior of *t*_div_ as a function of *θ*/*δ*_TE_ and In the divergent regime, after a few recombination events, the divergence *δ*(*t*) at the level of individual genes quickly escapes the integration barrier and increases indefinitely. Consequently, 〈Δ(*t*)〉 increases linearly with time (see e.g. =2 in Fig. 4 and Fig. 5) and reaches 〈Δ(*t*)〉 = 2*θ* within *∼*2*N*_*e*_ generations. In contrast for smaller values of *δ*_mut_/*δ*_TE_ in the metastable regime, it takes extremely long time for 〈Δ(*t*)〉 to reach 2*θ*. In this regime genes get repeatedly exchanged with the rest of the population and 〈Δ(*t*)〉 remains nearly constant over long periods of time (see e.g. *δ*_mut_/*δ*_TE_ = 2 × 10^−3^ in Fig. 4 and Fig. 5). Notably, near the boundary region between the two regimes a small perturbation in the evolutionary parameters could change the evolutionary dynamics from divergent to metastable and vice versa.

### Population structure

Can we understand the phylogenetic structure of the entire population by studying the evolutionary dynamics of just a single pair of strains?

Given sufficient amount of time every pair of genomes in our model would diverge indefinitely. However, in a finite population of size *N*_*e*_, the average probability of observing a pair of strains whose MRCA existed *t* generations ago is exponentially distributed, 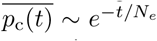 (here and below we use the bar to denote averaging over multiple realizations of the coalescent process, or longtime average over population dynamics) (27–29). Thus, while it may be possible for a pair of genomes considered above to diverge indefinitely from each other (see Fig. 4), it becomes more and more unlikely to find such a pair in a finite-sized population.

Let π (Δ) to denote the probability distribution of Δ for all pairs of genomes in a given population, while 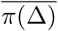 stands for the same distribution averaged over long time or multiple realizations of the population. One has

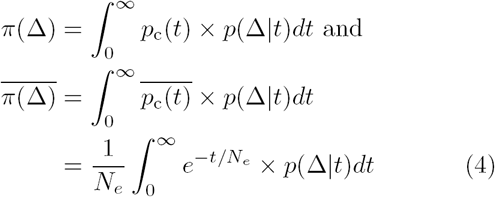

**FIG. 5.**
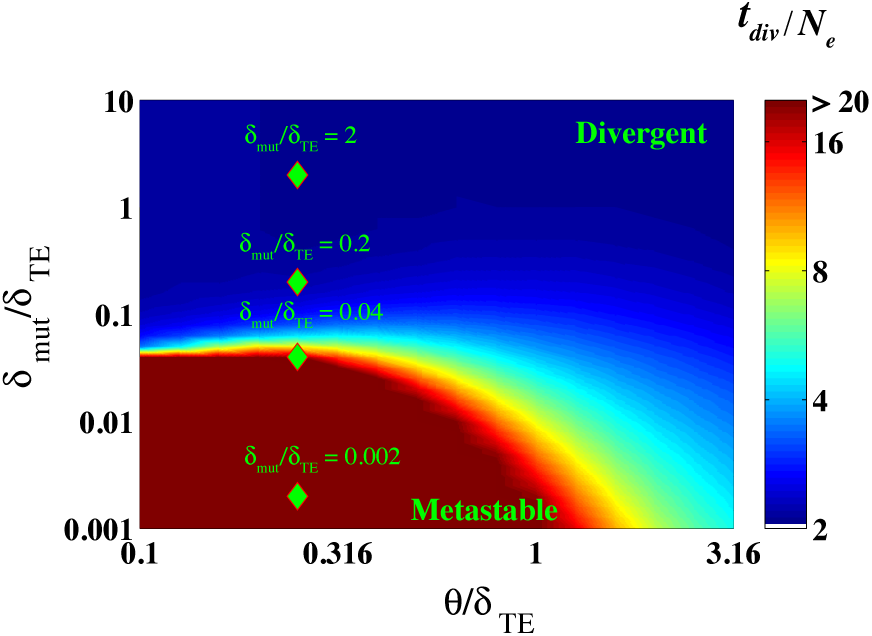
The number of generations *t*_div_ (in units of the population size *N*_*e*_) required for a pair of genomes to diverge well beyond the average intra-population diversity (see main text). We calculate the time it takes for the ensemble average 〈Δ(*t*)〉 of the genome-wide average divergence to reach 2*θ* as a function of *θ/δ*_TE_ and *δ*_mut_/*δ*_TE_. We used *δ*_TE_ = 1%, *μ* = 10^−2^ per gene per generation. In our simulations we varied *ρ* and *θ* to scan the (θ*δ*_TE_, *δ*_mut_/*δ*_TE_) space. The green diamonds represent four populations shown in Fig. 4 and Fig. 6 (see below).

In Eq. 4, *P*_c_ (*t*) is the probability that a pair of strains in the current population shared their MRCA *t* generations ago and *p*(Δ|*t*) is the probability that a pair of strains have diverged by Δ at time *t*. Given that Δ(*t*) is the average of G≫1 independent realizations of *δ*(*t*), we can approximate *p* (Δ|*t*) as a Gaussian distribution with average 〈*δ*(*t*)〉_G_ = ∫ *δ* × *p*(*δ*|*t*)*dδ* and vari ance 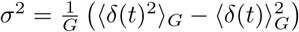. Here and below angular brackets and the subscript G denote the average of a quantity over the entire genome.

Unlike the time-or realization-averaged distribution 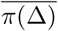, only the instantaneous distribution π (Δ) is accessible from genome sequences stored in databases. Notably, even for large populations these two distributions4 could be significantly different from each other. Indeed, *p*_*c*_(*t*) in any given population is extremely noisy due to multiple peaks from clonal subpopulations and does not resemble its smooth long-time average 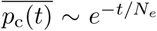 (28, 29). In panels a) to d) of Fig. 6, we show π (Δ) for the four cases shown in Fig. 4 (also marked by green diamonds in Fig. 5). We fixed the population size to *N*_*e*_ = 500. We changed *δ*_mut_/*δ*_TE_ by changing the recombination rate ρ. The solid lines represent a time snapshot obtained by numerically sampling *P*_*c*_ (*t*) in a Fisher-Wright population of size *N*_*e*_ = 500. The dashed black line represents the time average 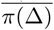.

**FIG. 6.**
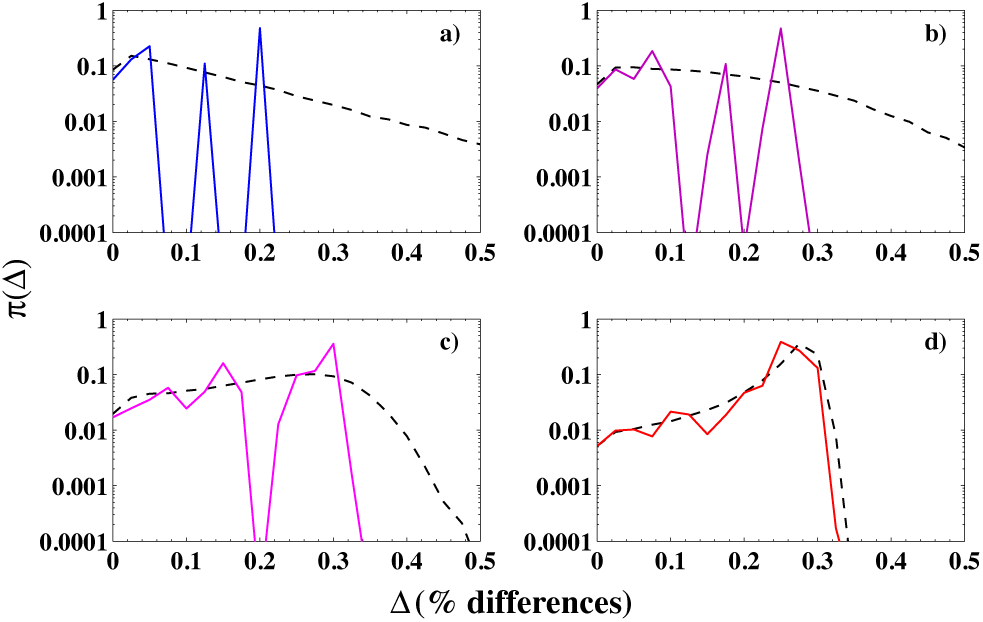
Distribution of all pairwise genome-wide divergences *δ*_*ij*_ in a co-evolving population for decreasing values of *δ*_mut_/*δ*_TE_: 2 in a), 0.2 in b), 0.04 in c) and 0.002 in d) In all 4 panels, dashed black lines represent time-averaged distributions 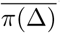, while solid lines represent typical “snap-shot” distributions π (Δ) in a single population. Colors of solid lines match those in Fig. 4 for the same values of parameters. Time-averaged and snapshot distributions were estimated by sampling 5 × 10^5^ pairwise coalescent times from the time-averaged coalescent distribution *p∼e*^*–t/N*^_*e*_ and the instantaneous coalescent distribution *p*_*c*_(*t*) correspondingly (see text for details).

In the divergent regime of Fig. 5, at high values of *δ*_mut_/*δ*_TE_ = μ/(*ρl*_*tr*_ *δ*_TE_), the instantaneous snapshot distribution π (Δ) has multiple peaks corresponding to divergence distances between several spontaneously formed clonal sub-populations present even in a homogeneous population. These sub-populations rarely exchange genetic material with each other, because of a low recombination frequency *ρ.* In this regime, the time averaged distribution 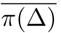 has a long exponential tail and, as expected, does not agree with the instantaneous distribution π (Δ).

Notably, in the metastable regime, at lower values of *δ*_mut_/*δ*_TE_, the exponential tail shrinks into a Gaussian-like peak. The width of this peak relates to fluctuations in Δ(*t*) around its mean value which in turn are dependent on the total number of genes *G*. Moreover, the difference between the instantaneous and the time averaged distributions decreases as well. In this limit, all strains in the population exchange genetic material with each other. Thus, in the metastable regime, frequent recombination events successfully eliminate multiple peaks due to clonal subs-populations thus forming a genetically cohesive and temporally stable population.

### Analysis of bacterial species

Where are real-life bacterial species located on the divergent-metastable diagram? Instead of *δ*_mut_/*δ*_TE_ as defined here, population genetic studies of bacteria usually quantify the relative strength of recombination over mutations as *r/m,* the ratio of the number of SNPs brought in by recombination relative to those generated by point mutations in a pair of closely related strains (6, 8, 12). In our framework, *r/m* is defined as *r/m* = *ρ*_succ_ /*μ* x *I*_tr_ x *δ*_tr_ where *ρ*_succ_ < *ρ* is the rate of successful recombination events and *δ*_tr_ is the average divergence in transferred regions. Both *ρ*_*succ*_ and *δ*_tr_ depend on the evolutionary parameters (see appendix for a detailed description of our calculations). *r/m* is closely related (but not equal) to the inverse of *δ*_mut_/*δ*_TE_ used in our previous plots.

In Fig. 7, we re-plotted the “phase diagram” shown in Fig. 5 in terms of *θ*/*δ*_TE_ and *r*/*m* and attempted to place several real-life bacterial species on it. To this end we estimated *θ* from the MLST data (30) and used *r/m* values that were determined previously by Vos and Didelot (12). As a first approximation, we assumed that the transfer efficiency *δ*_*TE*_ is the same for all species considered and is given by *δ*_*TE*_ *∼*2.26% used in Ref. (16). However, as mentioned above, the transfer efficiency *δ*_*TE*_ depends in part on the RM and the MMR systems. Given that these systems vary a great deal across bacterial species including minimal barriers to recombination observed e.g. in *Helicobacter pylori* (10) or different combinations of mul-. tiple RM systems reported in Ref. (31). We note that *Helicobacter pylori* appears divergent even with minimal barriers to recombination probably because of its ecologically structured population that is dependent on human migration patterns (32). One expects transfer efficiency *δ*_*TE*_ might also vary across bacteria. Further work is needed to collect the extent of this variation in a unified format and location. One possible bioinformatics strategy is to use the slope of the exponential tail in SNP distribution (*p*(*δ*|Δ) in our notation) to infer the transfer efficiency *δ*_*TE*_ as described in Ref. (6).

Fig. 7 allows one to draw the following conclusions., First, it confirms that both *r/m* and *θ*/*δ*_T E_ are important evolutionary parameters and suggests that each of them alone cannot fully classify a species as either divergent or metastable. Second, it predicts a sharp transition between the divergent and the metastable phases implying that a small change in *r/m* or *θ*/*δ*_TE_ can change the evolutionary fate of the species. And finally, one can, see that different bacterial species use diverse evolutionary strategies straddling the divide between these two regimes.

Can bacteria change their evolutionary fate? There are multiple biophysical and ecological processes by which bacterial species may move from the metastable to the divergent regime and vice versa in Fig. 5. For example, if the effective population size remains constant, a change in mutation rate changes both *δ*_mut_/*δ*_TE_ as well as *θ*. A change in the level of expression of the MMR genes, changes in types or presence of MMR, SOS, or restriction-modification (RM) systems, loss or gain of co-infecting phages, all could change *δ*_TE_ or the rate of recombination (14, 31) thus changing the placement of the species on the phase diagram shown in Fig. 7.

**FIG. 7.**
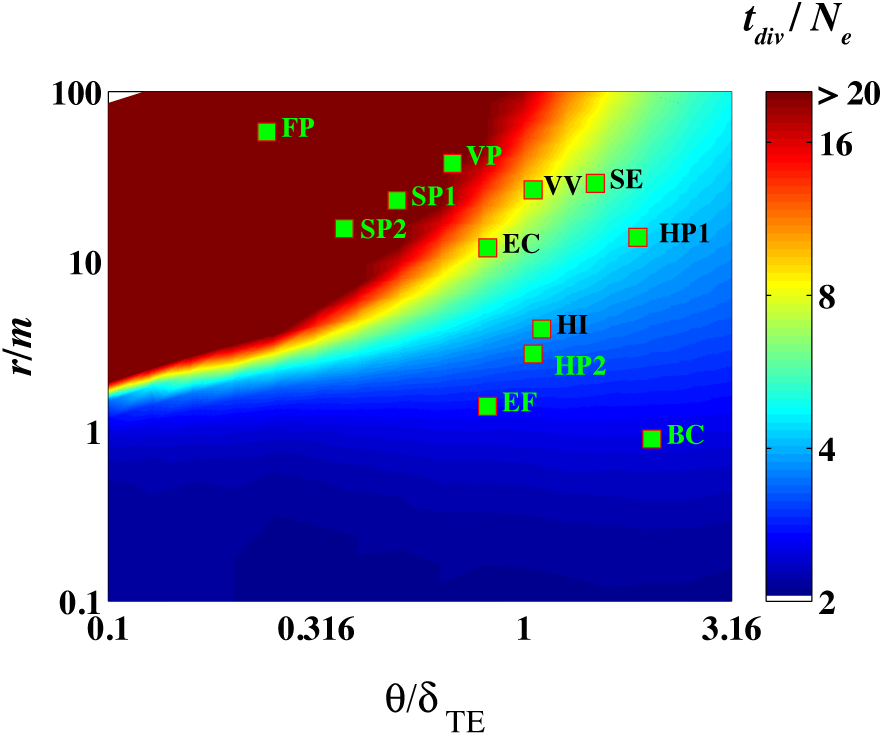
Approximate position of several real-life bacterial spaces on the metastable-divergent phase diagram (see text for details). Abbreviations of species names are as follows: FP: *Flavobacterium psychrophilum,* VP: *Vibrio para-haemolyticus,* SE: *Salmonella enterica,* VV: *Vibrio vulnificus,* SP1: *Streptococcus pneumoniae,* SP2: *Streptococcus pyogenes,* HP1: *Helicobacter Pylori,* HP2: *Haemophilus parasuis,* HI: *Haemophilus influenzae,* BC: *Bacillus cereus,* EF: *Enterococcus faecium,* and EC: *Escherichia coli.*

Adaptive and ecological events should be inferred from population genomics data only after rejecting the hypothesis of neutral evolution. However, the range of behaviors consistent with the neutral model of recombination-driven evolution of bacterial species was not entirely quantified up till now, leading to potentially unwarranted conclusions as illustrated in (33). Consider *E. coli* as an example. Known strains of *E. coli* are usually grouped into 5-6 different evolutionary sub-clades (groups A, B1, B2, E1, E2, and D). It is thought that inter-clade sexual exchange is lower compared to intra-clade exchange (6, 26). Ecological niche separation and/or selective advantages are usually implicated as initiators of such putative speciation events (17). In our previous analysis of 32 fully sequenced *E. coli* strains, we estimated *θ*/*δ*_TE_ > 3 and *r/m* ∼ 8 – 10 (6) implying that *E. coli* resides deeply in the divergent regime in Fig. 7. Thus, based on the analysis presented above one expects *E. coli* strains to spontaneously form transient sexually-isolated sub-populations even in the absence of selective pressures or ecological niche separation. In conclusion, a more careful analysis is needed to reject neutral models of evolution in the studies of population genetics of bacteria.

### EXTENDING THE FRAMEWORK

Throughout this study we used three main assumptions greatly simplifying the problem and allowing for exact mathematical analysis: i) exponentially decreasing probability of successful integration of foreign DNA into a recipient genome as a function of the local sequence divergence, ii) exponentially distributed pairwise coalescent time distribution of a neutrally evolving well-mixed population, and iii) independent evolution of non-overlapping “genes” or larger indivisible units of horizontal genetic transfer. Here we discuss how one can generalize the developed framework relax these assumptions.

(i) A wide variety of barriers to foreign DNA entry exist in bacteria (11). For example, *Helicobacter pylori*, is thought to have relatively free import of foreign DNA (10) while other bacteria may have multiple RM systems that either act in combination or are turned on and off randomly (31). Moreover, rare non-homologous/illegitimate recombination events can transfer highly diverged segments between genomes (11) potentially leading to homogenization of the population. Such events can be captured by a weaker-than-exponential dependence of the probability of successful integration on local genetic divergence (see Appendix for a calculation with non-exponential dependence of the probability of successful integration *p*_success_ on the local sequence divergence). One can incorporate these variations within our framework by appropriately modifying the functional relationship between the probability of successful integration and local sequence divergence or even by allowing it to change with time (e.g. relax recombination barriers in the presence of stress).

(ii) Bacteria belong to ecological niches defined by environmental factors such as availability of specific nutrient sources, host-bacterial interactions, and geographical characteristics. Bacteria in different environments may rarely compete with each other for resources and consequently may not belong to the same effective population and may have lowered frequency of DNA exchange compared to bacteria sharing the same niche. How can one capture the effect of ecological niches on genome evolution? Geographically and/or ecologically structured populations exhibit a coalescent structure (and thus a pair wise coalescence time distribution) that depends on the nature of niche separation (34, 35). Within our framework, niche-related effects can be incorporated by accounting for pairwise coalescent times of niche-structured populations (34, 35) and niche dependent recombination. frequencies. For example, one can consider a model with two or more subpopulations with different probabilities for intra-and inter-population DNA exchange describing geographical or phage-related barriers to recombination.

While most point mutations in bacterial genomes are thought to have insignificant fitness effect, the evolutionary dynamics of bacterial species is driven by rare advantageous mutations and thus is far from being neutral. Recombination in bacterial species is thought to be essential for their evolution in order to minimize the fitness loss due to Muller’s ratchet (36) and to minimize the impact of clonal interference (37). Thus, it is likely that both recombination frequency and transfer efficiency are under selection (36, 38, 39). How could one include fitness effects in our theoretical framework? Above, we considered the dynamics of neutrally evolving bacterial populations. The effective population size is incorporated in our framework only via the coalescent time distribution exp(–*T*/*N*_*e*_) and consequently the intra-species diversity exp(–*δ*/*θ*) (see supplementary materials). Neher and Hallatschek (40) recently showed that while pair-wise coalescent times in adaptive populations are not exactly exponentially distributed, this distribution has a pronounced exponential tail with an effective population size *N*_*e*_ weakly related to the actual census population size and largely determined by the variance of mutational fitness effects (40). In order to modify the recombination kernel *R*(*δ*_*a*_|*δ* _*b*_) one needs to know the 3-point coalescence distribution for strains *X*, *Y*, and the donor strain *D* (see Supplementary Materials here and in Ref. (6) for details). Once such 3-point coalescence distribution is available in either analytical or even numerical form our results could be straightforwardly generalized for adaptive populations (assuming most genes remain neutral). We expect the phase diagram of thus modified adaptive model to be similar to its neutral predecessor considered here, given that the pairwise coalescent time distribution in adaptive population has an exponential tail as well (40), and for our main results to remain qualitatively unchanged.

(iii) Finally, in this work, we assumed that recombination events transfer non-overlapping segments of length *l*_tr_ that always recombine in their entirety. In real bacteria, transfer events have variable lengths and partially overlap with each other (6, 9, 10, 41).

Do then the above conclusions about metastability in genome evolution hold when recombination tracks have variable end points and lengths? The metastability/divergent transition identified in this work (see Fig. 5 above) is based on the ensemble average 〈Δ(*t*)〉. We believe that while overlapping recombination events may play a role in determining correlations in local diversities *δ* along the chromosome, the the ensemble average of the pairwise divergence 〈Δ 〉 is likely to be insensitive to the nature of recombination.

We tested this hypothesis with an explicit simulation of *N*_*e*_ = 250 co-evolving strains each with *L*_*g*_ = 10^6^ base pairs. We performed three types of simulations (see panel

**FIG. 8.**
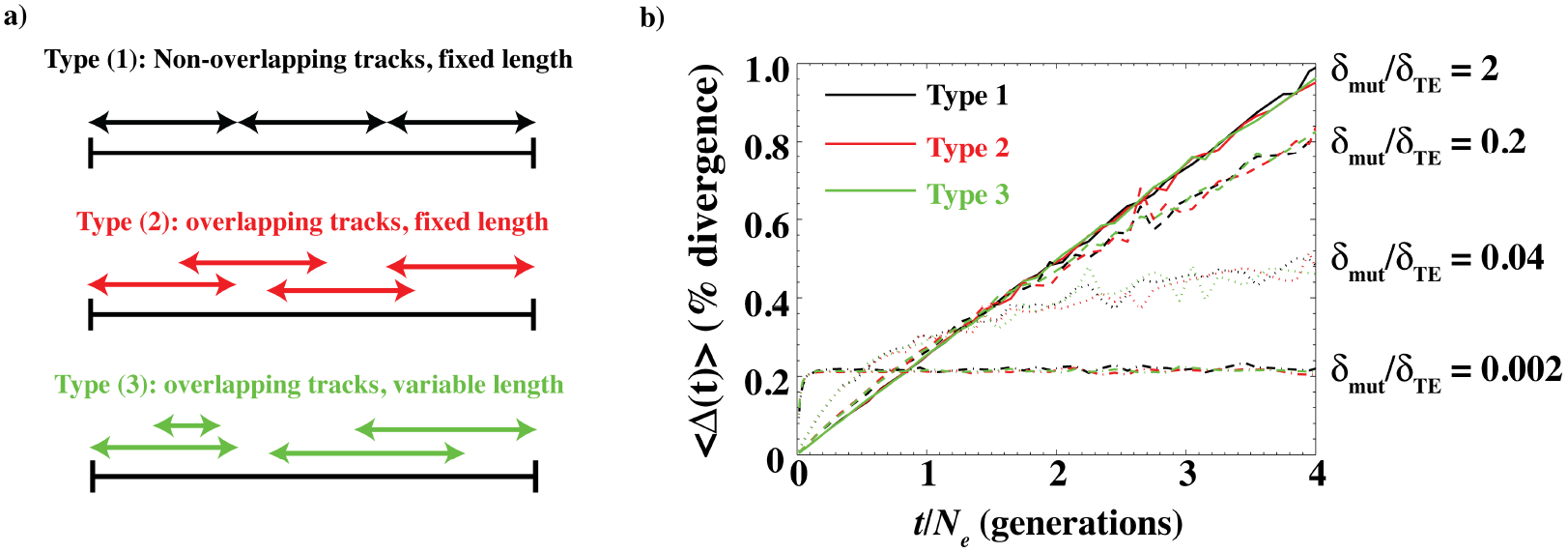
a) The schematics of the three types of simulations. In type (1), recombining stretches have fixed end points. As a result, different recombination tracks do not overlap. In type (2) and (3), the recombining stretches have variable end points and as a result different recombination tracks can potentially overlap with each other. b) The ensemble average ⟨Δ(*t*)⟩ of pairwise genome-wide divergence Δ(*t*) as a function of the pairwise coalescent time *t* in explicit simulations. Type (1) simulation has non-overlapping transfer of 5000 bp segments. Type (2) simulations have transfers of overlapping 5000 bp segments. Type (3) simulation have overlapping transfers of segments whose average length is 5000 bp. The value of *δ*_mut_/*δ*_TE_ is indicated on the top left corner.

We tested this hypothesis with an explicit simulation of Ne = 250 co-evolving strains each with *L*_*g*_ = 10^6^ base pairs. We performed three types of simulations (see panel a of Fig 8 for an illustration). In the first type (type (1)), every transfer event attempted a transfer of one genetic segment (of fixed length 5 kbp) in a non-overlapping manner. This protocol is identical to the one employed in this work. In the second type of simulation (type (2)), recombination tracks were allowed to start at any base pair but had a fixed length (of 5 kbp). Finally, we also, investigated the effect of variable track lengths. We ran a simulation (type (3)) where successful recombination events transferred on an average 5 kbp. The lengths of the recombination tracks were exponentially distributed with an average 5 kbp. We set the minimum transfer length to be 3 kbp. In order to directly compare results across different types of simulations, we ran each of the three simulations for the four parameter sets used in Fig. 4. See appendix for details of the simulations.

Panel b of Fig. 8 shows the time evolution of the ensemble average⟨Δ(*t*)) estimated from the explicit simulations. The three colors represent three different types of simulations. Notably, ⟨ Δ(*t*)) is insensitive to whether recombination tracks are of variable length or overlapping with each other. As mentioned above, the metastability explored in the manuscript is defined in terms of the ensemble average divergence⟨Δ(*t*)⟩. Consequently, we: believe that our quantitative and qualitative conclusions about metastability remain unchanged.

Can the effects of allowing overlapping recombination tracks be seen in population structure? Let us look at the stochastic fluctuations in Δ(*t*) around its ensemble average⟨Δ(*t*)). Intuitively, overlapping recombination events will homogenize highly divergent genetic fragments in the population. As a result, we expect smaller within population variation i.e. a smaller fluctuation in Δ(*t*) around⟨Δ(*t*)). We tested this by studying the numerical estimate of p(Δ) (see Eq. 4) for the three simulations.

We only consider the case where *δ*_mut_/ *δ*_TE_ = 0.002. As seen in Fig. 4 and Fig. 8, in the metastable state the divergence Δ(*t*) virtually does not increase as a function of *t* at long times (the rate of increase is extremely slow). Thus, the variance in 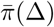 largely represents the variance in Δ(*t*) around its ensemble average ⟨Δ(*t*) ⟩. In Fig. 9, we show 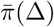 for the three different types of simulations. Notably, the variance in 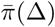 is much smaller when overlapping recombination events are allowed (type (2) and type (3) simulations compared to type (1) simulation). The effect of varying length of recombination events appears to be minimal. This suggests that variance in Δ(*t*) around its ensemble average⟨Δ(*t*)) is smaller when recombination tracks overlap with each other compared to a case where individual recombination events are independent of each other.

In short, while overlapping transfer events are likely to affect correlations in genetic diversity along the chromosome as well as the population structure, their role in determining the metastability/divergent transition described in this work appears minimal.

In our future studies we plan to explore these and other extensions on top of the basic mathematically tractable model described here.

## CONCLUSION

While recombination is now recognized as an important and sometimes even dominant contributor to patterns of genome diversity in many bacterial species(5, 6, 8–12), its effect on population structure and stability is still heavily debated (16, 17, 42–44). In this work, we explored three variants of a model of gene transfers in bacteria to study how the competition between mutations and recombinations affects genome evolution. Analysis of each of the three models showed that recombination-driven bacterial genome evolution can be understood as a balance between two important competing processes. We identified the two dimensionless parameters *θ*/*δ*_TE_ and *δ*_mut_/*δ*_TE_ that dictate this balance and result in two qualitatively different regimes in bacterial evolution, separated by a sharp transition.

**FIG. 9.**
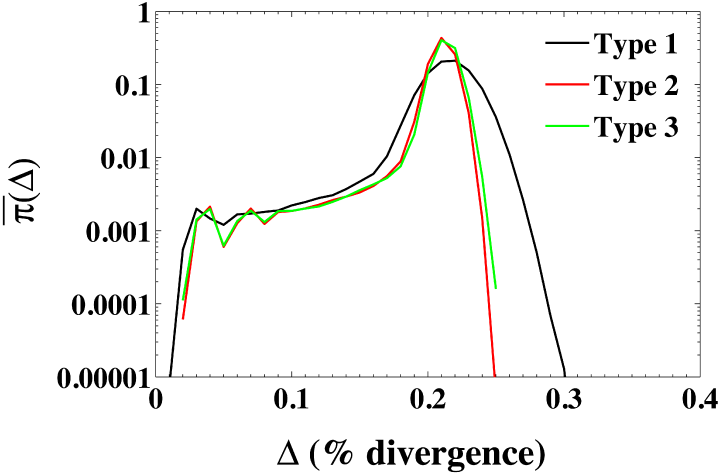
The ensemble average distribution of genome-wide divergence between pairs of strains 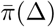 for the three types of simulations shown in panel a of Fig. 8 when *δ*_mut_/*δ*_TE_ = 0.002.

As seen in Fig. 5 and Fig. 7, in the divergent regime, the pull of recombination is insufficient to homogenize individual genes and entire genomes leading to a temporally unstable and sexually fragmented species. Notably, understanding the time course of divergence between a single pair of genomes allows us to study the structure of the entire population. As shown in Fig. 6, species in the divergent regime are characterized by multi-peaked clonal population structure. On the other hand, in the metastable regime, individual genomes repeatedly recombine genetic fragments with each other leading to a sexually cohesive and temporally stable population. As seen in Fig. 7, real bacterial species appear to belong to both of these regimes as well as in the cross-over region separating them from each other.

## Acknowledgments

We would like to thank Kim Sneppen, Erik van Nimwegen, Daniel Falush, Nigel Goldenfeld, Eugene Koonin, and Yuri Wolf for fruitful discussions and comments that lead to an improved manuscript. We would also like to thank the two reviewers and the editor for their detailed reading and valuable comments.

## APPENDIX

### ⟨Δ(*t*)⟩from computer simulations

We performed three types of explicit simulations of a Fisher-Wright population of N_*e*_ = 250 co-evolving strains. The three simulations had different modes of gene transfers as indicated in panel a of Fig. 8. Each strain had *L*_*g*_ = 10^6^ base pairs. Each base pair was represented either by a 0 (wild type) or 1 (mutated). The mutation rate was fixed at μ = 5 x 10^−6^ per base pair per generation. We varied the recombination rate ρ = 2.5 *x* 10^−8^, 2.5 *x* 10^−7^,1.25 *x* 10^−6^, and 2.5 *x* 10^−5^ per base pair per generation. *θ* was fixed at *θ* = 0.25% and *δ*_TE_ was fixed at *δ*_TE_ = 1%. These parameters are identical to the ones used in Fig. 4 of the main text. We note that given the low population diversity (*θ* = 0.25%), we can safely neglect back mutations.

Note that in all three types of simulations, on an average, a total of 5 kilobase pairs of genome was transferred in a successful transfer event thereby allowing us to directly compare the three simulations.

We strated the simulations with *N*_*e*_ identical genomes. We ran a Fisher-Wright simulation for 5000 = 20 x N_*e*_ generations to ensure that the population reached a steady state. In each generation, children chose their parents randomly. This ensured that the population size remained constant over time. Mutation and recombination events were attempted according to the corresponding rates. Note that it is non-trivial to keep track of the divergence between individual pairs over time since one or both of the strains in the pair may either be stochastically eliminated. To study the time evolution of the ensemble average⟨Δ(*t*)) of the divergence, at the end of the simulation, we collected the pairwise coalescent times *t* between all pairs of strains as well as Δ(*t*), the genomic divergences between them. Note that due to the stochastic nature of mutations and recombination events, Δ(*t*) is a random variable. We estimated the ensemble average⟨Δ(*t*) ⟩ by binning the pairwise coalescent times in intervals of dt = 25 generations (1/10th of the population size) and taking an average over all Δ(*t*) in each bin. The ensemble average thus estimated represents the average over multiple realizations of the coalescent process. Mathematically, the ensemble average is given by

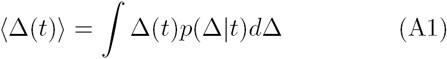

Here, *p*(Δ|*t*) is the probability that the genomes of two strains whose most recent common ancestor was *t* generations ago have diverged by Δ. We note that the variance in Δ(*t*) is expected to be small since it is an average over a large number of genes. We plot the ⟨Δ(*t*) ⟩ estimated from the three simulations in panel b) of Fig. 8.

**FIG. A1.**
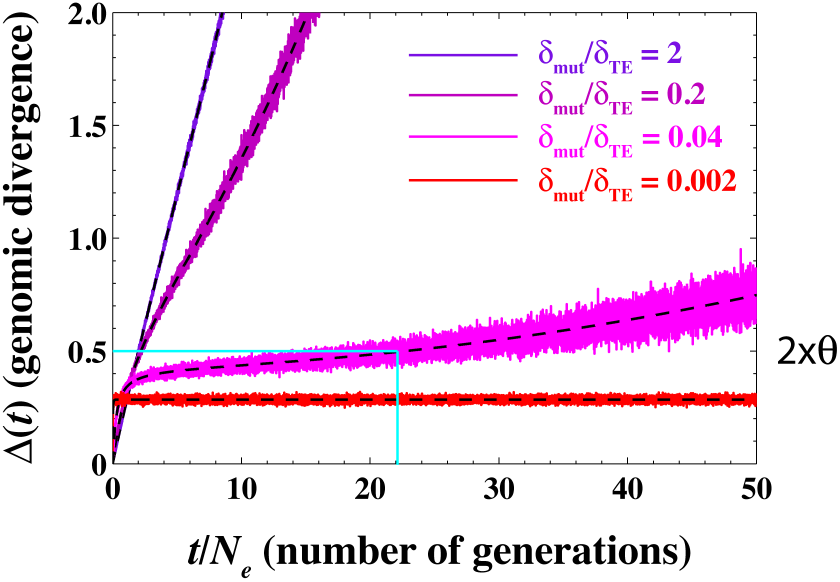
Genome-wide divergence Δ(*t*) as a function of time at *θ*/*δ*_TE_ = 0.25. We have used *δ*_TE_ = 1%, *θ* = 0.25%, *μ* = 10^−2^ per gene per generation and *ρ* = 10^−4^,10^−3^, 5 × 10^−2^, and 0.1 per gene per generation corresponding to *δ*_mut_/*δ*_TE_ = 2, 0.2, 0.04 and 2 x 10^−3^ respectively. The dashed black lines represent the ensemble average⟨Δ(*t*)⟩. The cyan lines show the time it takes for the ensemble-averaged genomic divergence ⟨ Δ(*t*)⟩ to reach 2*θ* when *δ*_mut_/*δ*_TE_ = 0.04 (pink line).

### Behavior of ⟨Δ(t)⟩ in the long time limit

### Estimating *r/m*

As mentioned in the main text, *r*/*m* is defined in a pair of strains as the ratio of SNPs brought in by recombination events and the SNPs brought in by point mutations. Clearly, *r*/*m* will depend on a strain-to-strain comparison however, usually it is reported as an average over all pairs of strains. How do we compute *r*/*m* in our framework? We have

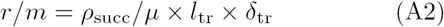

Thus, in order to compute *r*/*m*, we need two quantities. First, we need to compute the rate of successful recombinations *ρ*_sub>succ/sub>_ ρ. We can calculate *ρ*_succ_ as

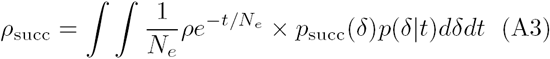

where psucc is the success probability that a gene that has diverged by *δ* will have a successful recombination event. The integration over exponentially distributed pairwise coalescent times averages over the population. psucc can be computed from Eq. 3 by integrating over all possible scenarios of successful recombinations. We have

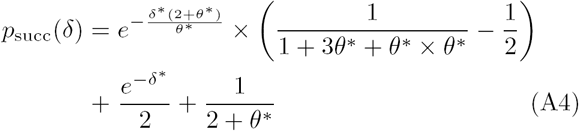

where *δ*\* = *δ*/*δ*_TE_ and *θ*\* = *θ*/*δ*_TE_ are normalized divergences and p(*δ*|*t*) is the distribution of local divergences at time *t*. In practice, *r*/*m* can only estimated by analyzing statistics of distribution of SNPs on the genomes of closely related strain pairs where both clonally inherited and recombined parts of the genome can be identified (6, 26). Here, we limit the time-integration in Eq. A3 to times *t* < min(N_e_ = *θ*/2*μ*, *δ*_TE_/2*μ*).

Second, we need to compute the average divergence in transferred segments, *δ*_*tr*_. We have

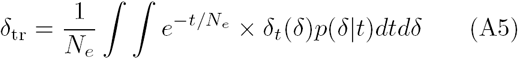

where *δt*(*δ*) is the average divergence after a recombination event if the divergence before transfer was *δ*.

### Computing *θ* from MLST data

Except for *E. coli* where we used our previous analysis (6) (we used *θ*/*δ*_TE_ ∼3 and *r*/*m* = 12), we downloaded MLST sequences of multiple organisms from the MLST database (30). For each of the 7 genes present in the MLST database, we performed a pairwise alignment between strains. *θ* for each gene was calculated as the average of pairwise SNPs. The *θ* for the species was estimated as average of the *θ*_s_ calculated for each of the 7 genes.

### Non-exponential dependence of *P*_success_ on local sequence divergence

In the main text, we showed that when Psucess decays exponentially with the local divergence, the time evolution of local divergence *δ*(*t*) shows metastability. When the recombination rate is low, a few recombination events take place that change *δ*(*t*) to typical values in the population before the local region eventually escapes the integration barrier, leading to a linear increase in *δ*(*t*) (see Fig. 3). When the recombination rate is high, the number of recombination events before the eventual escape from the integration barrier increases drastically leading to metastable behavior.

Here, we suggests that weaker-than-exponential dependence of *P*_success_ can lead to a time evolution of local divergence *δ*(*t*) that never escapes the integration barrier, leading to a genetically homogeneous population independent of the recombination rate *ρ*.

While it is difficult to carry out analytical calculations for a finite *θ* and *δ*_TE_, following Doroghazi and Buckley (20), we consider the limit *θ* → 0 when *μ* and *ρ* are finite. The time evolution of *δ*(*t*) in the limit *θ* → 0 when *P*_success_ decays exponentially with divergence is given by (see Eq. 3)

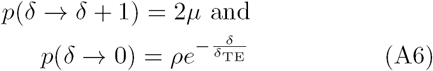

In Eq. A6, *δ*(*t*) is the number of SNPs (as opposed to SNP density used in the main text). As was shown in the main text, the evolution of *δ*(*t*) described by Eq. A6 is a random walk that repeatedly resets to zero before eventually escaping to *δ* → ∞. The number of resetting events depends on *δ*_mut_/*δ*_TE_ as defined in the main text (see low *θ*/*δ*_TE_ values in Fig. 5).

A generalization to non-exponential dependence of the success probability is straightforward,

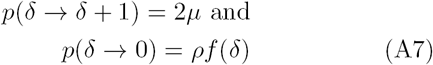

where 1 ≤ *f*(*δ*) ≥ 0 is the probability of successful integration. How weak should the integration barrier *f*(*δ*) be so that the time evolution described by Eq. A7 can never escape the pull of recombination? In other words, whatc are the conditions on *f*(*δ*) that ensure that the time evolution of local divergence described by Eq. A7 results inc a random walk that resets to zero infinitely many times?

If the random walk resets infinitely many times, it hasc a well defined stationary distribution as *t* → ∞.Notec that the random walk described by an exponentially de-caying Psuccess does not have a well defined stationaryc distribution since as *t* → ∞, *δ*(*t*) → ∞ regardless of the rate of recombination and the transfer efficiency. Let us assume that f (*δ*) is such that there exists a well-defined stationary distribution. We define *P*_*i*_as the probability that *δ* = *i* in the stationary state. We can write balance equations in the stationary state

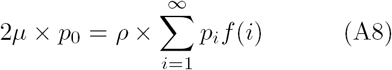

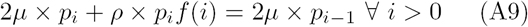

Rearranging

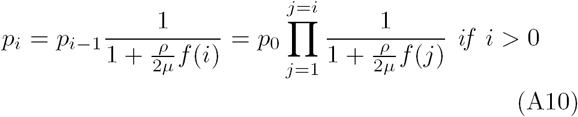

Since *P*_0_ ≠ 0, from Eq. A9 and Eq. A10 we have for an arbitrary *f*(*δ*) (denoting *ρ*/2*μ* = *t*)

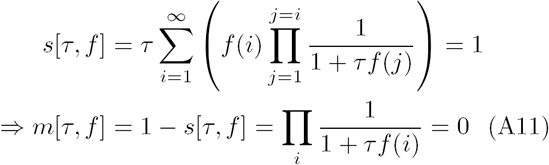

Thus, as long as the *functional s*[*τ*, *f*] in Eq. A11 is equal to 1 (or *m*[*τ*, *f*] = 0), the walk remains localized. Eq. A11 is a surprisingly simple result and is valid for any 0 ≤ *f*(*δ*) ≤ 1.

Let us consider a specific case where *f*(*δ*) = *δ*^−*ν*^. A power-law dependence in *P*_success_ is weaker than the exponential decay used in the main text, potentially allowing transfers between distant bacteria. Let us examine the self-consistency condition. We have

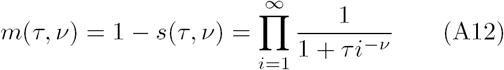

Taking logarithms and using the Abel-Plana formula

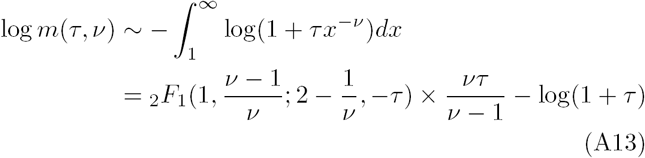

if *ν* ≥ 1. The integral (and thus the sum) tends to *∞* when *ν* < 1. Here, _2_*F*_1_ is the hypergeometric function. Thus, when *ν* < 1, a well defined stationary distribution exists and as long as *ρ* > 0 and *μ* > 0 regardless of ρ and the population remains genetically cohesive. When *ν* > 1, we expect behavior similar to the exponential case studied in the main text, viz. a divergent vs metastable transition depending on the competition between forces of recombinations and mutations. We believe that these conclusions will also hold true when *θ* is finite.

